# Intrinsic T-cell programming and immune spatial organization govern sex-biased tuberculosis immunity

**DOI:** 10.64898/2026.07.07.737119

**Authors:** Manish Gupta, Stefanie Krug, Sarita Neupane, Moagi Shaku, Sabal Chaulagain, Shichun Lun, Joseph P. Hoffmann, Eileen Scully, Sabra L. Klein, William R. Bishai

## Abstract

Biological sex can profoundly influence the susceptibility to infectious diseases, yet the mechanisms behind the sex-dependent protective immunity against tuberculosis (TB) remain poorly understood. Here we show that sexually divergent immunity during chronic *Mycobacterium tuberculosis* (Mtb) infection is governed by both intrinsic T cell programming and pulmonary immune spatial organization. Using the Four Core Genotype (FCG) mouse model, adoptive cell transfer, pathway-specific blockade and B cell depletion, we demonstrate that CD4⁺ T cells from gonadal females (XXF), but not XX males (XXM), confer enhanced protection to susceptible XY male recipients, independently of sex chromosome complement. Female-derived CD4⁺ T cells reduce Mtb burdens while promoting pulmonary Bcl6⁺ CD4⁺ T cell responses and limiting neutrophilic inflammation. Mechanistically, blockade of CXCR3 or CD40L abrogates female-associated protection, with CD40L signaling additionally required to maintain organized pulmonary B cell structures. Although depletion of conventional B-2 B cells did not impair bacterial control, it disrupted tertiary lymphoid organization and revealed striking sex-specific functions of pulmonary B cells. Loss of B cell follicles (BCFs) primarily remodeled adaptive T cell responses in females, whereas in males it drove inflammatory myeloid activation, exaggerated neutrophil recruitment and widespread neutrophil extracellular trap (NET) formation. Together, these findings identify two complementary layers of sex-dependent immune regulation during TB: intrinsic programming of protective female CD4⁺ T cells, and B cell-dependent spatial organization that coordinates adaptive immunity in females while restraining pathological inflammation in males. These findings establish immune tissue organization as a key determinant of the sexually dimorphic host defense during chronic TB.

## Introduction

TB remains one of the leading infectious causes of mortality worldwide and exhibits marked sex-associated differences in disease susceptibility and outcome. Globally, males develop active pulmonary TB more frequently than females and experience greater disease severity, mortality, and treatment failure^1, 2^. Experimental and clinical studies indicate that biological sex profoundly influences host immunity, with females generally mounting stronger innate and adaptive immune responses, whereas males develop exaggerated inflammatory pathology during chronic Mtb infection^3, 4, 5, 6^. Although sex steroid hormones have been implicated in these differences, the cellular mechanisms by which biological sex shapes protective immunity and disease progression during TB remain poorly understood.

Protective immunity against Mtb depends primarily on adaptive immune responses, particularly IFN-γ-producing CD4⁺ T cells^7^, with CD8⁺ T cells also contributing importantly to bacterial control during chronic infection^8, 9^. Beyond conventional Th1 responses, specialized CD4⁺ T cell subsets, including CXCR5⁺ Bcl6⁺ follicular helper T (Tfh)-like cells, coordinate local immune responses by promoting B cell activation, germinal center formation, and tertiary lymphoid structure (TLS) development within infected lungs^10, 11, 12^. Sex steroid hormones regulate both T cell differentiation and Tfh responses^3, 13^, raising the possibility that intrinsic sex-dependent programming of adaptive immunity contributes to the divergent TB outcomes observed in males and females.

Growing evidence further implicates the spatial organization of the pulmonary immune response as a critical determinant of TB outcome^14^. In both human disease and experimental models, pulmonary BCFs and TLSs are associated with improved bacterial control, enhanced local antigen presentation, and coordinated adaptive immune responses while limiting excessive inflammation^12, 15, 16^. Rather than functioning solely through antibody production, B cells regulate immunity through antibody-independent mechanisms that include antigen presentation, cytokine production, and organization of local immune microenvironments^16, 17^. Disruption of B cell compartments alters granuloma architecture, enhances inflammatory pathology, and perturbs T cell activation during chronic infection^18, 19, 20, 21^. Maintenance of these organized immune structures depends upon coordinated interactions between T cells, B cells, and chemokine networks that regulate immune cell positioning within infected tissues. Among these pathways, CXCR3 has emerged as an important regulator of leukocyte trafficking and granuloma organization^22, 23^, whereas CD40L-dependent interactions are essential for T cell–B cell communication and lymphoid tissue formation^24, 25^. However, whether these pathways contribute to sex-specific immune organization and protection during chronic TB remains unknown.

Using the Four Core Genotype (FCG) mouse model to genetically separate gonadal and chromosomal sex^26^, we previously identified gonadal sex, rather than sex chromosome complement, as the principal determinant of susceptibility to chronic TB^27^. Both XX and XY gonadal males developed increased bacterial burden, excessive neutrophilic inflammation, disrupted granuloma organization, and enhanced NET formation, whereas XX gonadal females exhibited superior bacterial control associated with coordinated T and B cell responses and prominent pulmonary B cell follicle formation^27^. These findings indicate that biological sex influences not only immune cell composition but also the spatial organization of adaptive immunity within infected lungs^27^. However, it remains to be determined if female-biased protection reflects intrinsic differences in adaptive immune cell functions, whether these responses regulate pulmonary immune architecture, and how organized lymphoid structures contribute to protection and immunopathology during chronic infection.

Here, we combined adoptive cell transfer, antibody-mediated pathway blockades and B-cell depletion in the FCG model to further define the mechanisms underlying female-biased protection during chronic TB. We demonstrate that CD4⁺ T cells from gonadal females, but not XX males, transfer enhanced protection independently of sex chromosome complement, identifying gonadal sex as the dominant determinant of protective CD4⁺ T cell programming. We further identify CXCR3- and CD40L-dependent pathways as key regulators of protective adaptive immunity and pulmonary lymphoid organization. Finally, we show that pulmonary B cells primarily function as organizers of immune spatial architecture rather than direct mediators of bacterial control, with strikingly different roles in the two sexes: in females, B-2 B cells maintain organized adaptive immune niches, whereas in males, they restrain inflammatory myeloid remodeling, neutrophil recruitment, and NET-associated pathology. Together, our findings establish a mechanistic framework in which sex-dependent immunity during TB is governed by two complementary layers of regulation: intrinsic programming of adaptive immune cells and B cell-dependent organization of pulmonary immune architecture.

## Results

### XX female, but not XX male-derived CD4⁺ T cells confer protection against Mtb infection in XY males

Our previous studies using the FCG mouse model identified a striking dichotomy between TB-susceptible gonadal males, which develop excessive myeloid inflammation and disrupted granuloma organization, and TB-resistant XX gonadal females, which exhibit coordinated adaptive immune responses and enhanced pulmonary B cell follicle formation during chronic TB. To determine whether sex-specific programming of adaptive immune cells contributes directly to these divergent outcomes, we performed adoptive transfer experiments using congenic XX female (XXF; CD45.1) and XX male (XXM; CD45.2) donor mice, thereby isolating the effects of gonadal sex independent of sex chromosome complement. CD4⁺ T cells or CD19⁺ B cells from Mtb-infected donors were transferred into chronically infected XYM recipients (Fig. 1a).

**Figure 1.**
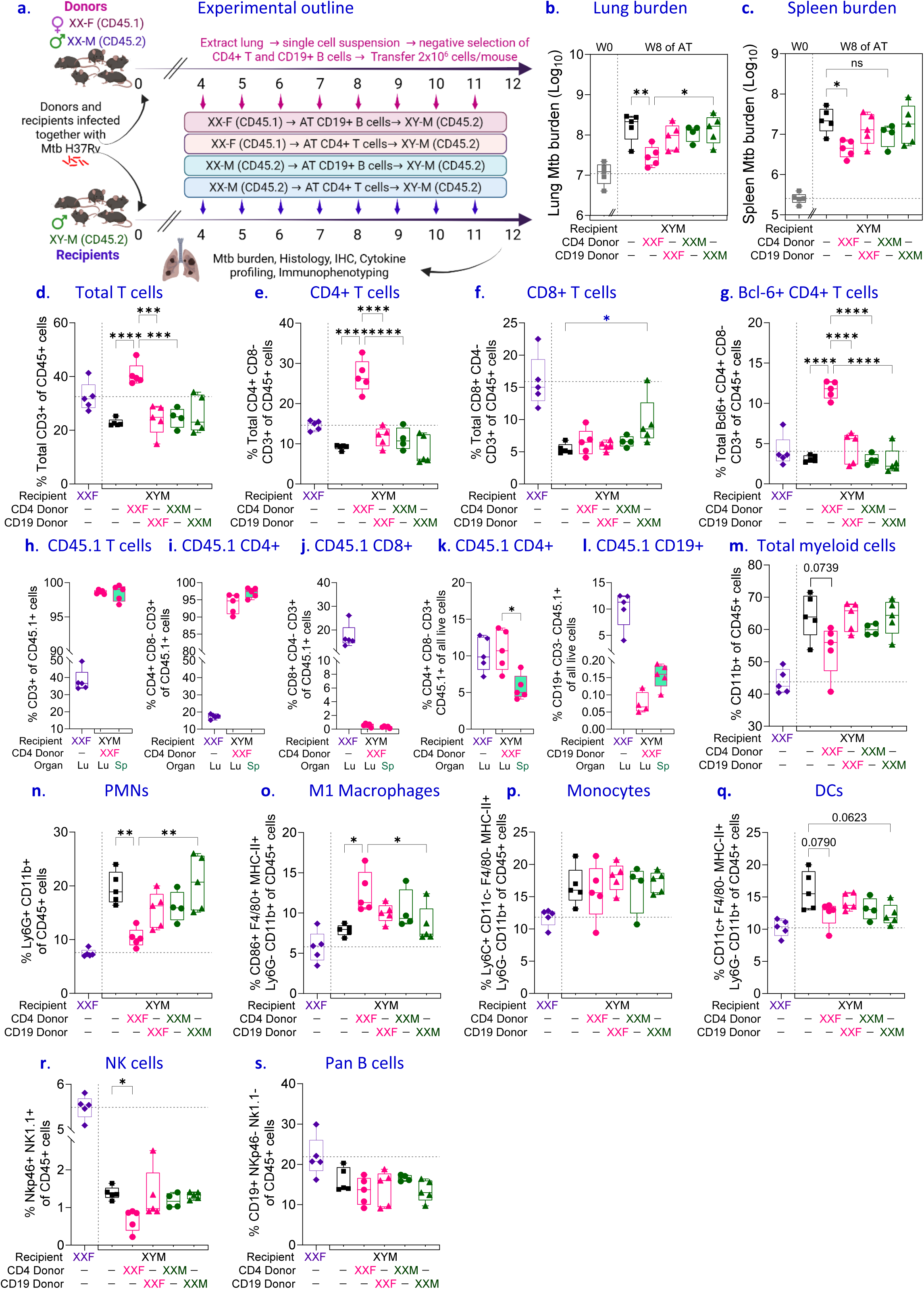
XXF-derived CD4⁺ T cells confer enhanced protection against Mtb infection and promote Bcl6⁺ CD4⁺ T-cell responses in XYM recipients. **(a)** Experimental design. Female (XXF; CD45.1) and male (XXM; CD45.2) donor mice were infected with *Mtb* H37Rv together with XYM recipient mice (CD45.2). Four weeks post-infection (wpi), 2 × 10⁶ CD4⁺ T cells or CD19⁺ B cells were isolated from donor lungs and adoptively transferred weekly into Mtb-infected XYM recipients. Recipient mice were analyzed at 12 weeks post-infection (wpi), after eight consecutive adoptive transfers, for bacterial burden and immune cell composition. (c) Lung and (c) spleen Mtb burden at 12 wpi. W0 denotes bacterial burden in recipient mice at 4 wpi, prior to adoptive transfer (grey bars). Recipients of XXF-derived CD4⁺ T cells exhibited reduced pulmonary and splenic bacterial burden compared with recipients of XXM-derived CD4⁺ T cells. (d–g) Flow cytometric analysis of pulmonary lymphocyte populations. (d) total CD3⁺ T cells, (e) CD4⁺ T cells, (f) CD8⁺ T cells, and (g) Bcl6⁺ CD4⁺ T cells among CD45⁺ lung leukocytes. Transfer of XXF-derived CD4⁺ T cells increased total CD4⁺ T-cell and Bcl6⁺ CD4⁺ T-cell frequencies relative to other groups. (h–l) Characterization of donor-derived CD45.1⁺ cells in recipient tissues. **(h)** frequency of donor-derived CD45.1⁺ cells among total CD45⁺ cells in lung and spleen, **(i)** donor-derived CD4⁺ T cells, **(j)** donor-derived CD8⁺ T cells, **(k)** donor-derived CD4⁺ T cells among live CD45.1⁺ cells, and **(l)** donor-derived CD19⁺ B cells among live CD45.1⁺ cells. XXF donor CD4⁺ T cells persisted and accumulated in recipient tissues following transfer. **(m–s)** Analysis of pulmonary innate and lymphoid immune populations. **(m)** total myeloid cells, **(n)** neutrophils (PMNs), **(o)** M1 macrophages, **(p)** monocytes, **(q)** dendritic cells, **(r)** NK cells, and **(s)** pan B cells among CD45⁺ lung leukocytes. Data are presented as box-and-whisker plots showing individual mice. n = 5 for all groups except XYM receiving XXM-derived CD4⁺ T cells, where n = 4. Boxes represent the interquartile range, and whiskers denote minimum and maximum values. Statistical significance was determined using one-way ANOVA with Tukey’s multiple comparisons test. *P < 0.05, **P < 0.01, ***P < 0.001, ****P < 0.0001, and ns, not significant.

Transfer of XXF-derived CD4⁺ T cells resulted in a marked improvement of Mtb infection control in XYM recipients, indicated by significantly reduced bacterial burdens in both the lungs and spleens compared with infected XYM males that did not receive an adoptive transfer (Fig. 1b,c). This effect was unique to XXF-derived CD4+ T cells, as neither CD4+ T cells from XXM nor CD19+ B cells from either XXF or XXM donors significantly impacted bacterial burdens of XYM recipients. To define the immune mechanisms associated with this protection, pulmonary lymphocyte populations were analyzed by flow cytometry. The enhanced protection in XXF CD4⁺ T cell-recipient XYM mice correlated with the striking accumulation of total CD3⁺ T cells within the lungs, driven predominantly by expansion of the CD4⁺ T-cell compartment (Fig. 1d,e). In contrast, the transfer of XXM-derived CD4+ T cells did not affect CD3+ or CD4+ T cell frequencies in XYM recipients, and CD8⁺ T cell frequencies remained largely unchanged among all treatment groups (Fig. 1f). Notably, transfer of XXF CD4⁺ T cells was associated with a significant enrichment of Bcl6⁺ CD4⁺ T cells (Fig. 1g), suggesting enhanced development or maintenance of T follicular helper (Tfh)-like responses. Given the established role of Bcl6-expressing CD4⁺ T cells in promoting local immune organization and protective immunity during TB infection, these findings suggest that XXF-derived CD4⁺ T cells preferentially support adaptive immune programs associated with improved disease control, while XXM-derived CD4^+^ T cells may exhibit reduced pulmonary homing capacity. Analysis of XXF-derived cells confirmed successful engraftment and persistence of transferred CD45.1⁺ cells in both the lungs and spleens of recipient animals (Fig. 1h). The XXF-derived CD45.1+ population was predominantly composed of CD4⁺ T cells, with minimal representation of CD8⁺ T cells (Fig. 1i,j). Furthermore, transferred XXF CD4⁺ T cells homed and accumulated efficiently within the XYM recipient lung, resulting in a comparable CD45.1+ frequency to XXF recipients (Fig. 1k), supporting their direct role in facilitating protection.

Previous studies have shown that the adoptive transfer of splenic CD19⁺ B cells from Mtb-infected mice into B cell-deficient mice can rescue enhanced morbidity and aberrant granulomatous responses following aerosol Mtb infection despite limited tissue infiltration, with few donor-derived B cells detected in the spleen and virtually none in the lungs^21^. Therefore, to maximize the transfer of Mtb-specific B cells with potential for pulmonary homing, CD19⁺ B cells (>90% purity) were isolated from the lungs of chronically infected XXF and XXM donors. However, consistent with previous reports, and unlike CD4⁺ T cells, repeated transfer of lung-derived B cells resulted in only minute populations of XXF-derived CD45.1⁺ B cells detectable in recipient lungs and spleens at the experimental endpoint (Fig. 1l), indicative either of limited survival, failed lung/spleen homing, trafficking bias or niche dependency of transferred B cells. Accordingly, CD19⁺ B cell transfer from either XXF or XXM donors had no measurable effect on Mtb burden (Fig. 1b,c); however, the poor recovery of donor B cells precludes definitive conclusions regarding their contribution to host resistance in this setting.

Interestingly, only the transfer of XXF but not XXM CD4⁺ T cells altered the pulmonary immune landscape in XYM recipients. Although the frequencies of total CD11b⁺ myeloid cells and CD11c+ dendritic cells (DCs) were only modestly decreased (Fig. 1m,q), XXF CD4+ T cell recipients displayed evidence of broader immune remodeling characterized by significantly decreased neutrophil infiltration and increased M1 macrophage frequencies (Fig. 1n,o), while monocyte populations remained largely unchanged (Fig. 1p). NK cell frequencies were also reduced only in XYM mice receiving XXF-derived CD4⁺ T cells (Fig. 1r). Total pulmonary B cell frequencies remained relatively stable across experimental groups (Fig. 1s), indicating that any observed protection did not appear to be driven by major alterations in the B cell compartment.

Collectively, these findings demonstrate that the transfer of specifically XXF-derived CD4⁺ T cells is sufficient to enhance the resistance to chronic TB in susceptible XYM hosts, associated with increased bacterial containment and the expansion of CD4⁺ T cell responses, particularly Bcl6⁺ Tfh-like cells. The differential protective capacities of XXF and XXM donor CD4⁺ T cells despite an identical XX chromosome complement indicate that gonadal sex, rather than sex chromosome complement, is the dominant determinant of protective CD4⁺ T cell programming during Mtb infection.

### XXF-derived CD4⁺ T cells reduce pulmonary immunopathology and inflammatory responses during chronic Mtb infection in XYM recipients

To determine whether the enhanced bacterial control mediated by XXF-derived CD4⁺ T cells was accompanied by altered lung pathology, histological and inflammatory analyses were performed on lungs collected from recipient mice at the experimental endpoint (Fig. 1a). Histopathological examination of H&E-stained lung sections revealed marked differences in pulmonary inflammation between groups (Fig. 2a). XYM recipients receiving XXF CD4⁺ T cells displayed reduced inflammatory infiltration and increased preservation of alveolar architecture compared with recipients of XXM CD4⁺ T cells. Quantitative histopathological scoring confirmed a significant reduction in overall lung involvement following the transfer of XXF-derived CD4⁺ T cells (Fig. 2b). Because perivascular and peribronchiolar leukocyte cuffing are hallmarks of TB-associated immunopathology, these parameters were also evaluated. Recipients of XXF CD4⁺ T cells displayed significantly lower peribronchiolar leukocyte cuff (PBLC) and perivascular leukocyte cuff (PVLC) scores than mice receiving XXM donor CD4⁺ T cells (Fig. 2b,c), indicating diminished pulmonary inflammation. Since CD19⁺ B cell transfer did not alter bacterial burdens or immune cell accumulation (Fig. 1) and no overt differences in pulmonary histopathology or inflammatory responses were observed (Fig. 2a), these findings were not examined further.

**Figure 2.**
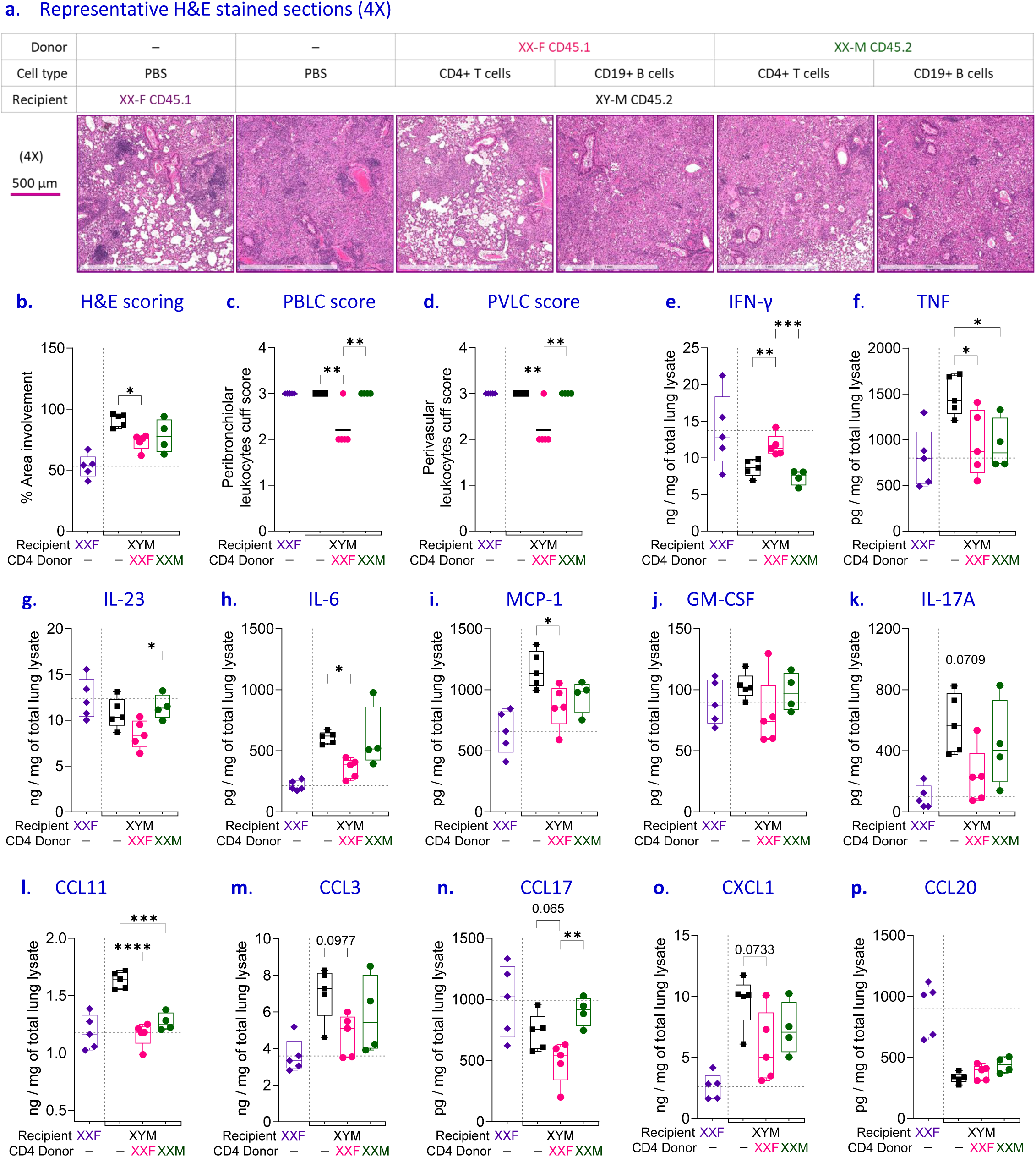
XXF-derived CD4⁺ T cells reduce pulmonary immunopathology and modulate inflammatory cytokine and chemokine responses during chronic Mtb infection in XYM recipients. **(a)** Representative H&E-stained lung sections (4× magnification) from recipient mice at the experimental endpoint. **(b–d)** Quantification of lung pathology. **(b)** percentage of total lung area involved by inflammatory lesions, **(c)** peribronchiolar leukocyte cuff (PBLC) score, and **(d)** perivascular leukocyte cuff (PVLC) score. Recipients of female-derived CD4⁺ T cells exhibited reduced pulmonary inflammation and leukocyte cuffing compared with recipients of male-derived CD4⁺ T cells. **(e–k)** Cytokine concentrations measured in lung lysates. **(e)** IFN-γ, **(f)** TNF, **(g)** IL-23, **(h)** IL-6, **(i)** MCP-1, **(j)** GM-CSF, and **(k)** IL-17A. **(l–p)** Chemokine concentrations in lung lysates. **(l)** CCL11 (Eotaxin), **(m)** CCL3, **(n)** CCL17, **(o)** CXCL1, and **(p)** CCL20. Data are presented as box-and-whisker plots showing individual mice. n = 5 for all groups except XYM receiving XXM-derived CD4⁺ T cells, where n = 4. Boxes represent the interquartile range and whiskers denote minimum and maximum values. Statistical significance was determined using one-way ANOVA with Tukey’s multiple comparisons test. *P < 0.05, **P < 0.01, ***P < 0.001, ****P < 0.0001, and ns, not significant.

To further characterize the pulmonary inflammatory milieu, cytokine concentrations were quantified in lung lysates. Transfer of XXF CD4⁺ T cells was associated with significantly higher IFN-γ (Fig. 2e) and lower IL-23 and IL-6 (Fig. 2g,h) levels than transfer of XXM CD4⁺ T cells, whereas TNF concentrations were reduced in response to XXF- or XXM-derived CD4+ T cells (Fig. 2f). In contrast, GM-CSF levels were unchanged across groups (Fig. 2j). IL-17A showed a trend toward lower expression in recipients of XXF donor cells, although this difference did not reach statistical significance (Fig. 2k). Chemokine profiling further revealed attenuation of inflammatory signals in XYM mice receiving female-derived CD4⁺ T cells. While both the macrophage/monocyte/neutrophil chemoattractant CCL3 and the major neutrophil-recruiting chemokine CXCL1 were trending toward reduced levels following XXF- but not XXM-derived CD4+ T cell transfer (Fig. 1m,o), lung levels of eotaxin/CCL11 were reduced in response to either XXF or XXM donor cells (Fig. 2l). XXF- or XXM-derived CD4⁺ T cells had no effect on lung CCL20 levels, which are predominantly elevated in females (Fig. 2p). Collectively, these findings indicate that female-derived CD4⁺ T cells confer protection against chronic Mtb infection by reducing bacterial burden while limiting pulmonary inflammation and tissue pathology, identifying intrinsic sex-dependent differences in CD4⁺ T cell function as key regulators of host resistance and disease outcome during TB.

### CXCR3- and CD40L-dependent pathways regulate pulmonary immune organization during chronic Mtb infection in females

Our previous study using the FCG model demonstrated that females exhibit superior control of chronic Mtb infection and identified *Cxcr3* and *Cd40lg* among the most highly differentially expressed immune genes in gonadal females relative to gonadal males at both 4 and 12 weeks post-infection (wpi). Both genes were consistently enriched in females by RNA sequencing and independently validated by qPCR (Fig. 3a–d). Our subsequent adoptive transfer studies established female-derived CD4⁺ T cells as the principal mediators of TB resistance, implicating sex-biased T cell programs in the sexually divergent susceptibility to TB. Because CXCR3 and CD40L are predominantly expressed by activated antigen-experienced T cells—including Th1, T follicular helper (Tfh), and memory CD4⁺ T cell populations, with CXCR3 also marking activated CD8⁺ T cells during chronic TB—we hypothesized that these pathways contribute to the female-specific immune landscape associated with protection. Accordingly, Mtb-infected XXF mice were treated with neutralizing antibodies against CXCR3 or CD40L from 4 to 12 weeks post infection, followed by assessment of bacterial burden, pulmonary pathology, immune cell composition, and B-cell organization at the experimental endpoint (Fig. 3e).

**Figure 3.**
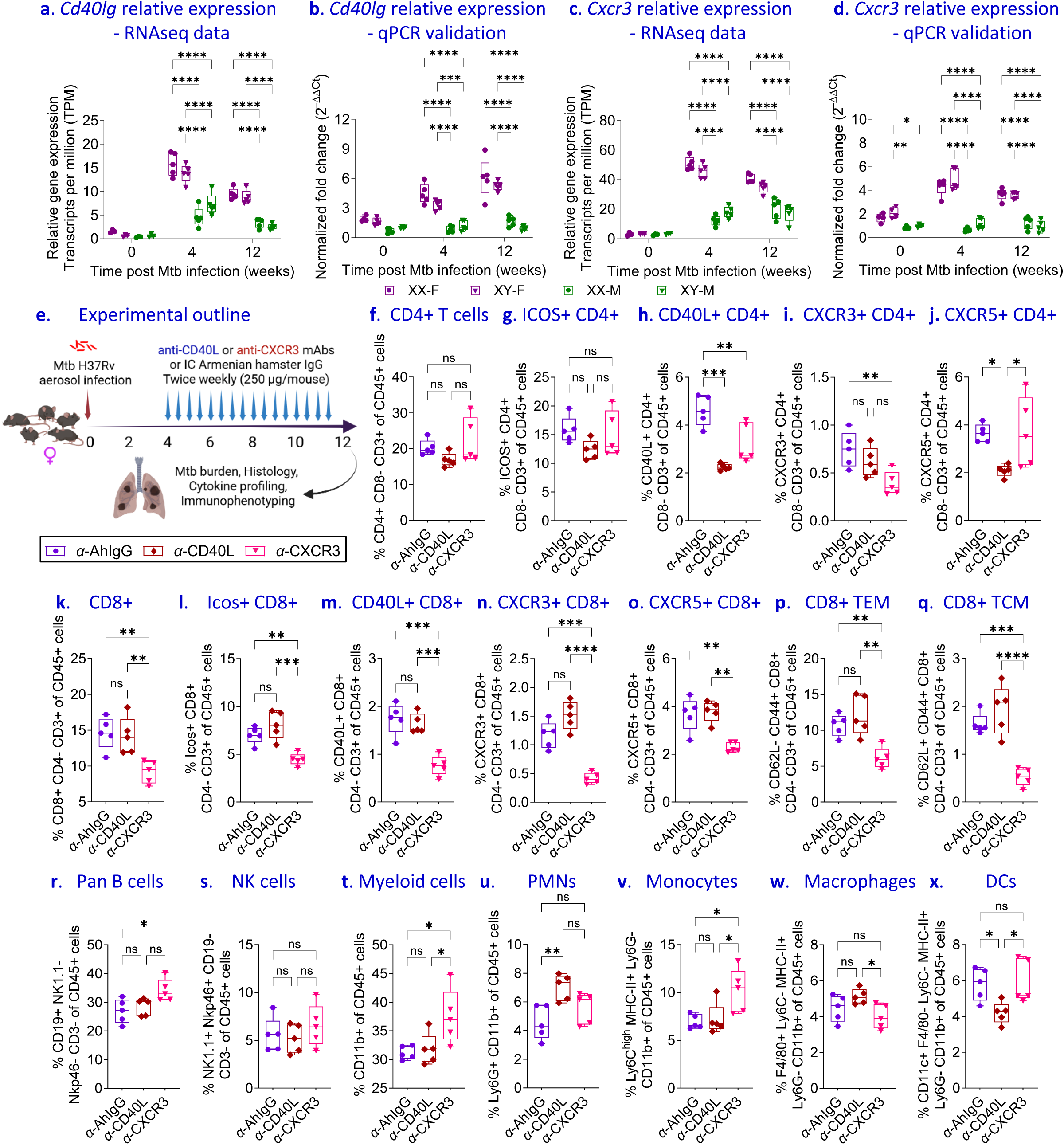
Female-biased CXCR3 and CD40L expression identifies immune pathways regulating pulmonary leukocyte composition during Mtb infection. **(a–d)**, Differential expression of *Cxcr3* and *Cd40lg* in the lungs of XX- and XY-gonadal female and male FCG mice at 0, 4, and 12 weeks post-Mtb infection was quantified. **(a, c)**, transcripts per million (TPM) counts obtained from RNA sequencing. **(b, d)**, qPCR validation of pulmonary *Cxcr3* and *Cd40lg* expression. **(e)**, Experimental design. XX female mice were infected with Mtb and treated biweekly with isotype control, anti-CXCR3, or anti-CD40L antibodies from 4 to 12 wpi. Lungs and spleen were harvested at 12 wpi for immunophenotyping and CFU enumeration. Flow cytometric analysis of pulmonary immune cell populations, including **(f)** total CD4⁺ T cells, **(g)** ICOS⁺ CD4⁺ T cells, **(h)** CD40L⁺ CD4⁺ T cells, **(i)** CXCR3⁺ CD4⁺ T cells, **(j)** CXCR5⁺ CD4⁺ T cells, **(k)** total CD8⁺ T cells, **(l)** ICOS⁺ CD8⁺ T cells, **(m)** CD40L⁺ CD8⁺ T cells, **(n)** CXCR3⁺ CD8⁺ T cells, **(o)** CXCR5⁺ CD8⁺ T cells, **(p)** effector memory (TEM; CD44⁺CD62L⁻) CD8⁺ T cells, **(q)** central memory (TCM; CD44⁺CD62L⁺) CD8⁺ T cells, **(r)** B cells, **(s)** NK1.1⁺ NKp46⁺ NK cells, **(t)** CD11b⁺ myeloid cells, **(u)** PMNs, **(v)** inflammatory monocytes, **(w)** macrophages, and **(x)** dendritic cells. Data are presented as box-and-whisker plots. n = 5 for all groups, and each symbol represents an individual mouse. Data are plotted as mean ± SEM. Statistical significance was determined using one-way ANOVA with Tukey’s multiple comparisons test. *P < 0.05, **P < 0.01, ***P < 0.001, ****P < 0.0001, and ns, not significant.

Despite antibody-mediated blockade, the overall frequency of pulmonary CD4⁺ T cells and ICOS⁺ CD4⁺ T cells remained largely unchanged (Fig. 3f,g). As expected, anti-CD40L treatment markedly reduced CD40L⁺ and CXCR5⁺ CD4⁺ T cell frequencies, confirming effective target engagement (Fig. 3h,j). In contrast, CXCR3 blockade selectively remodeled the CD4⁺ T cell compartment by reducing the minor CXCR3⁺ CD4⁺ T cell population, whereas CXCR5⁺ CD4⁺ T cells remained unchanged (Fig. 3i,j).

The effects of CXCR3 inhibition were more pronounced within the CD8⁺ T cell compartment. Total CD8⁺ T cells, together with ICOS⁺, CD40L⁺, CXCR3⁺, and CXCR5⁺ CD8⁺ T-cell populations, were all reduced following CXCR3 blockade (Fig. 3k–o). Likewise, both effector memory (TEM) and central memory (TCM) CD8⁺ T cell subsets were diminished (Fig. 3p,q), demonstrating a broad requirement for CXCR3 signaling in maintaining pulmonary CD8⁺ T cell responses during chronic Mtb infection.

Perturbation of these T cell-associated pathways also induced extensive remodeling of the pulmonary innate immune compartment. CXCR3 blockade increased pulmonary B cell frequencies without affecting NK cells (Fig. 3r,s). Total CD11b⁺ myeloid cells were expanded, accompanied by increased monocyte frequencies, whereas macrophages, neutrophils and dendritic cells were unaffected (Fig. 3t–x). In contrast, anti-CD40L treatment increased PMN and reduced DC frequencies while B cells, NK cells, total myeloid cells and monocytes remained unchanged (Fig. 3r-x). Together, these findings indicate that female-enriched CXCR3- and CD40L-dependent immune programs regulate pulmonary leukocyte organization during chronic Mtb infection, but that disruption of CXCR3- and CD40L-dependent signaling extends beyond T cell responses to reshape the broader immune landscape of the chronically Mtb-infected female lung. Although these interventions primarily target activated T cell populations, they induce widespread remodeling of both adaptive and innate immune compartments, providing the foundation for determining how these pathways influence bacterial control and disease outcome.

### Blockade of CXCR3 or CD40L abrogates female T cell-associated protection during chronic Mtb infection

To further evaluate the contribution of CXCR3- and CD40L-dependent immune programs to the female-associated protection against chronic TB, bacterial burden, lung pathology, B cell organization and inflammatory mediators were assessed in Mtb-infected female (XXF) mice following antibody-mediated blockade (Fig. 4). Indeed, neutralization of either CXCR3 or CD40L significantly impaired bacterial control, resulting in increased bacterial burdens in both the lungs and spleens at 12 weeks post infection compared with isotype-treated controls (Fig. 4a,b). Histopathological evaluation revealed an unexpected divergence between bacterial burden and tissue pathology. Despite significantly increased lung bacterial burdens following CXCR3 or CD40L blockade, quantitative analysis of H&E-stained lung sections demonstrated reduced histopathology scores, whereas disease severity indices remained comparable across treatment groups (Fig. 4c-e). Because organized B cell aggregates constitute a prominent component of pulmonary inflammatory lesions in protected females, we hypothesized that the observed reduced lung involvement reflected the disruption of lymphoid organization rather than attenuation of inflammation. Consistent with this interpretation, pathological assessment of alveolar macrophages and polymorphonuclear neutrophils (PMNs) revealed no significant differences among treatment groups (Fig. 4f,g), while representative H&E sections demonstrated reduced lymphoid organization, particularly following CD40L blockade (Fig. 4e).

**Figure 4.**
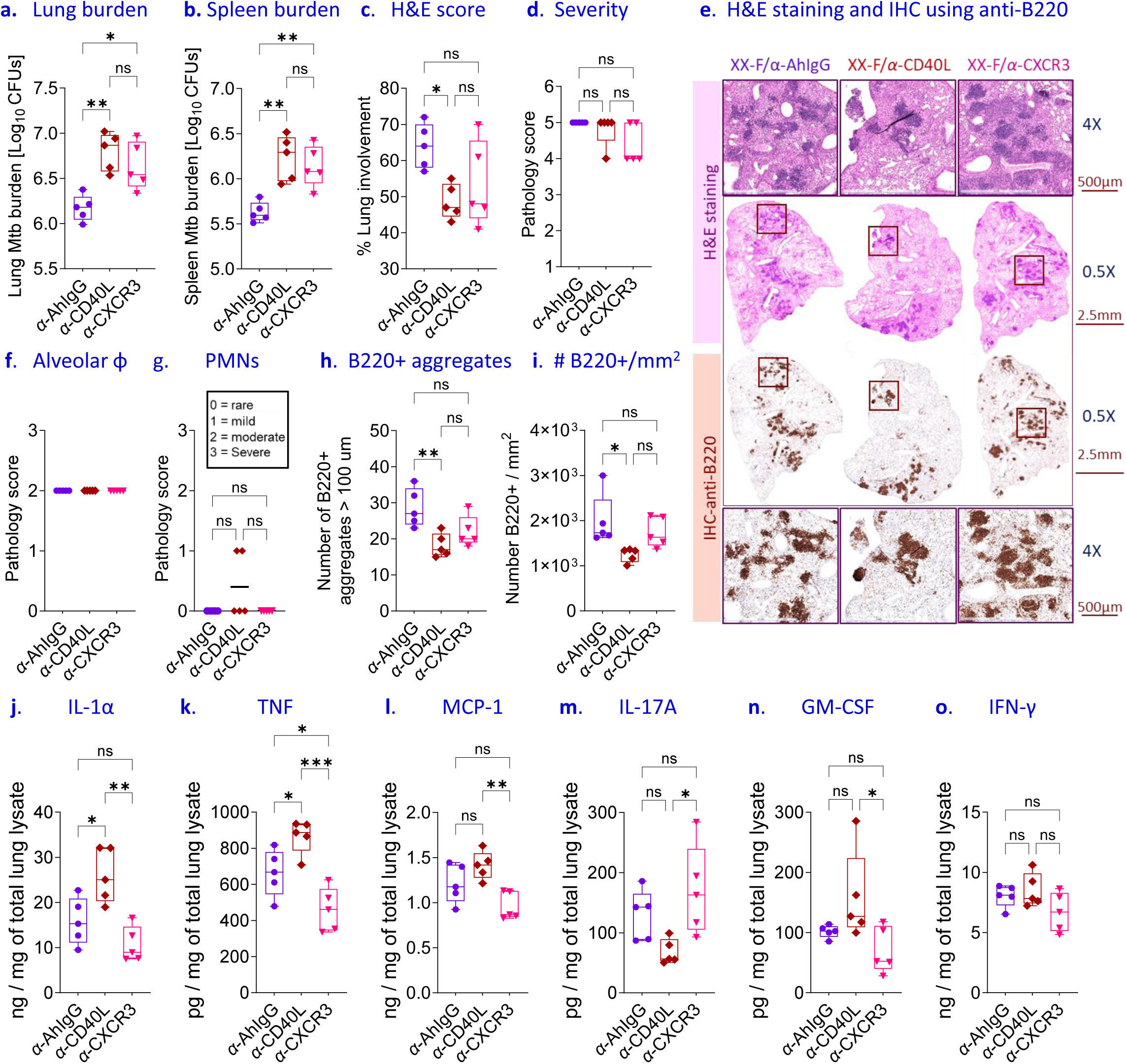
Blockade of CXCR3 or CD40L impairs bacterial control and alters pulmonary immune organization during chronic Mtb infection. Pulmonary and systemic bacterial burdens in XXF mice treated bi-weekly with isotype control, anti-CXCR3, or anti-CD40L antibodies from 4 to 12 weeks post-infection were enumerated. **(a)**, Lung, and **(b)**, spleen Mtb burden at 12 weeks post infection. Histopathological analysis of lung sections showing **(c)**, histopathology scores, **(d)**, disease severity scores, and **(e)**, representative hematoxylin and eosin (H&E)-stained lung sections together with representative B220 immunohistochemistry. Quantification of inflammatory cell pathology: **(f)** alveolar macrophage scores and **(g)** PMN scores. QuPath analysis of pulmonary B-cell organization showing **(h)**, number of B220⁺ lymphoid aggregates > 100μm, and **(i)**, B220⁺ cell density per mm^2^. Lung cytokine concentrations determined by LEGENDplex immunoassay showing-**(j)** IL-1α, **(k)** TNF, **(l)** MCP-1 (CCL2), **(m)** IL-17A, **(n)** GM-CSF, and **(o)** IFN-γ. Data are presented as box-and-whisker plots. n = 5 for all groups, and each symbol represents an individual mouse. Data are plotted as mean ± SEM. Statistical significance was determined using one-way ANOVA with Tukey’s multiple comparisons test. *P < 0.05, **P < 0.01, ***P < 0.001, ****P < 0.0001, and ns, not significant.

Given the established association between female TB resistance and pulmonary B cell aggregate formation, lung sections were next examined for B220⁺ lymphoid structures. CD40L blockade profoundly disrupted pulmonary B cell organization, significantly reducing both the number of B220⁺ aggregates and the density of B220⁺ cells within the lung parenchyma compared with isotype-treated controls (Fig. 4h,i). Representative anti-B220 immunohistochemistry confirmed a marked loss of organized B cell structures following CD40L neutralization (Fig. 4e). CXCR3 blockade also showed a trend toward reduced B220⁺ aggregate formation, although these changes did not reach statistical significance (Fig. 4h,i). Together, these findings support the notion that the lower histopathology scores observed following CD40L blockade primarily reflect disruption of organized B cell-rich inflammatory structures rather than reduced pulmonary inflammation.

To further characterize the inflammatory milieu associated with pathway inhibition, cytokine concentrations were quantified in lung homogenates. CD40L blockade increased IL-1α concentrations relative to isotype-treated controls, whereas CXCR3 blockade had no appreciable effect (Fig. 4j). Consistent with increased PMN infiltration, TNF concentrations were elevated following CD40L blockade but reduced after CXCR3 inhibition compared to control groups (Fig. 4k). MCP-1 concentrations were trending toward being reduced following CXCR3 blockade and elevated after CD40L neutralization but neither of these trends were significant (Fig. 4l). In contrast, IL-17A concentrations were non-significantly reduced following CD40L blockade but increased after CXCR3 inhibition (Fig. 4m). GM-CSF and IFN-γ concentrations remained largely unchanged across the treatment groups (Fig. 4n,o). Altogether, these findings demonstrate that both CXCR3- and CD40L-dependent pathways are required for female CD4⁺ T cell-mediated protection during chronic Mtb infection. Although blockade of either pathway impaired bacterial control, CD40L neutralization additionally disrupted pulmonary B cell organization, resulting in the loss of organized B220⁺ aggregates despite increased bacterial burden. Together, these data identify CXCR3 and CD40L as complementary regulators of protective immunity and pulmonary immune architecture in females during chronic TB.

### Pulmonary B cells maintain sex-specific immune spatial organization and differentially restrain inflammatory remodeling during chronic TB

Because adoptive transfer of lung-derived Mtb-specific B cells resulted in poor donor cell persistence and failed to definitively establish their contribution to host resistance, we next examined the function of endogenous pulmonary B cells by depleting CD20⁺ B cells during established chronic Mtb infection (Fig. 5a). Due to the complexity of these experiments, we restricted our initial experiments solely to 2 of the four FCG types, i.e. the two most dissimilar in B cell follicle formation, namely XXF (highest BCFs) vs. XYM (lowest BCFs). Histopathological analysis demonstrated that B cell depletion altered pulmonary immune organization differentially in XXF and XYM mice. Although overall lung involvement was only modestly affected (Fig. 5b,c), depletion of CD20⁺ B cells resulted in an almost complete loss of organized B220⁺ lymphoid aggregates throughout the lungs of XXF mice (Fig. 5d). In contrast, XYM lungs, which contained relatively sparse B cell structures at baseline, exhibited little organized lymphoid architecture following anti-CD20 treatment, confirming that continuous B cell maintenance is required to sustain the prominent tertiary lymphoid structures that characterize the resistant female lung.

**Figure 5.**
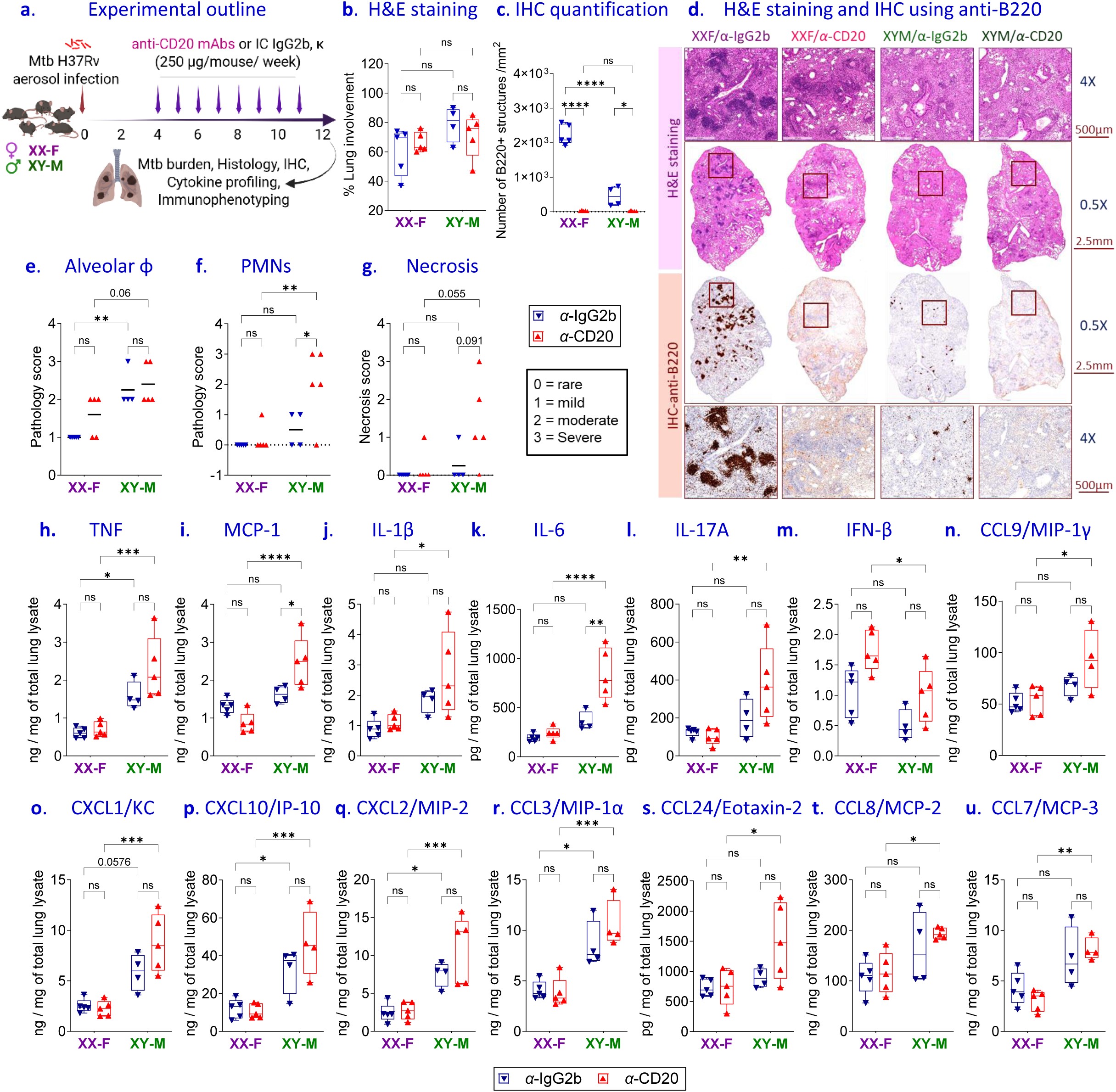
Conventional B-cell depletion disrupts sex-specific spatial organization of pulmonary immunity and differentially remodels inflammatory responses during chronic Mtb infection. **(a),** Experimental design depicting anti-CD20 mAb or IgG2b isotype control treatment of XXF and XYM mice from 4 to 12 weeks after aerosol Mtb infection. **(b),** Histopathological analysis of hematoxylin and eosin (H&E)-stained lung sections. **(c),** Quantification of anti-B220 immunohistochemical staining. **(d),** Representative H&E-stained lung sections together with representative anti-B220 immunohistochemistry depicting pulmonary B-cell organization in isotype control- and anti-CD20-treated XXF and XYM mice. **(e),** Alveolar macrophage pathology scores. **(f),** PMN pathology scores. **(g),** Pulmonary necrosis scores. Lung cytokine and chemokine concentrations determined by LEGENDplex immunoassay showing-**(h)** TNF, **(i)** MCP-1 (CCL2), **(j)** IL-1β, **(k)** IL-6, **(l)** IL-17A, **(m)** IFN-β, **(n)** CCL9/MIP-1γ, **(o)** CXCL1/KC, **(p)** CXCL10/IP-10, **(q)** CXCL2/MIP-2, **(r)** CCL3/MIP-1α, **(s)** CCL24/Eotaxin-2, **(t)** CCL8/MCP-2, and **(u)** CCL7/MCP-3. Data are presented as box-and-whisker plots. n = 5 for all groups except for IgG2b isotype-control-treated XYM, where n=4, and each symbol represents an individual mouse. Data are plotted as mean ± SEM. Statistical significance was determined using two-way ANOVA with Tukey’s multiple comparisons test. *P < 0.05, **P < 0.01, ***P < 0.001, ****P < 0.0001, and ns, not significant.

Despite comparable effects on B cell depletion, the pathological consequences diverged markedly between the sexes. Anti-CD20 treatment produced only modest alterations in alveolar macrophage accumulation in female or male mice (Fig. 5e). While PMN pathology and necrosis similarly unaffected in XXF mice, XYM mice exhibited greater neutrophilic infiltration accompanied by a trend toward increased pulmonary necrosis (Fig. 5f–g). Thus, while B cell depletion disrupted pulmonary immune architecture in both sexes, loss of these organized structures preferentially promoted inflammatory tissue remodeling in male mice.

To determine whether disruption of pulmonary B cell organization altered the local inflammatory milieu, cytokines and chemokines were quantified in lung homogenates. B cell depletion elicited only relatively modest cytokine changes in XXF mice but induced broad inflammatory activation in XYM mice. Concentrations of TNF, MCP-1 (CCL2), IL-1β, IL-6, IL-17A, IFN-β and CCL9/MIP-1γ were consistently increased following anti-CD20 treatment in males (Fig. 5h–n), indicating amplification of inflammatory pathways associated with myeloid-cell activation. Likewise, multiple leukocyte-recruiting chemokines, including CXCL1/KC, CXCL10/IP-10, CXCL2/MIP-2, CCL3/MIP-1α and CCL24/Eotaxin-2, were markedly elevated in B cell-depleted XYM mice (Fig. 5o–s), whereas CCL8/MCP-2 and CCL7/MCP-3 were comparatively less affected (Fig. 5t,u). The preferential induction of neutrophil-attracting chemokines suggested that loss of pulmonary B cell organization predominantly unleashes inflammatory myeloid recruitment in males. Collectively, these findings demonstrate that pulmonary B cells are essential for maintaining the spatial organization of the chronically infected lung but exert fundamentally different functions in the two sexes. In females, B cells sustain the organized B220⁺ lymphoid structures that typify protective pulmonary immunity, whereas in males they primarily restrain inflammatory cytokine and chemokine networks. These observations suggest that disruption of pulmonary immune architecture does not simply eliminate B cells but instead initiates sexually divergent immune programs that differentially shape chronic TB pathology.

### Disruption of pulmonary B-2 B cell organization differentially remodels adaptive and innate immune landscapes in females and males during chronic TB

Having established that anti-CD20 treatment disrupted pulmonary B cell-rich lymphoid structures and differentially altered inflammatory pathology (Fig. 5), we next investigated how the loss of these organized B cell niches reshapes the pulmonary immune landscape during chronic Mtb infection. Flow cytometric analysis confirmed efficient depletion of conventional B cell populations responsible for pulmonary follicle formation (Fig. 6a). Anti-CD20 immunotherapy resulted in near-complete ablation of B2 and follicular B (FoB) cells in both XXF and XYM mice (Fig. 6b-c), consistent with the profound loss of B220⁺ lymphoid aggregates observed histologically (Fig. 5d). In contrast, B1 B cell frequencies were minimally affected (Fig. 6d). These findings confirm that anti-CD20 selectively targets conventional B cell subsets that maintain pulmonary lymphoid organization while largely sparing the B1 compartment.

**Figure 6.**
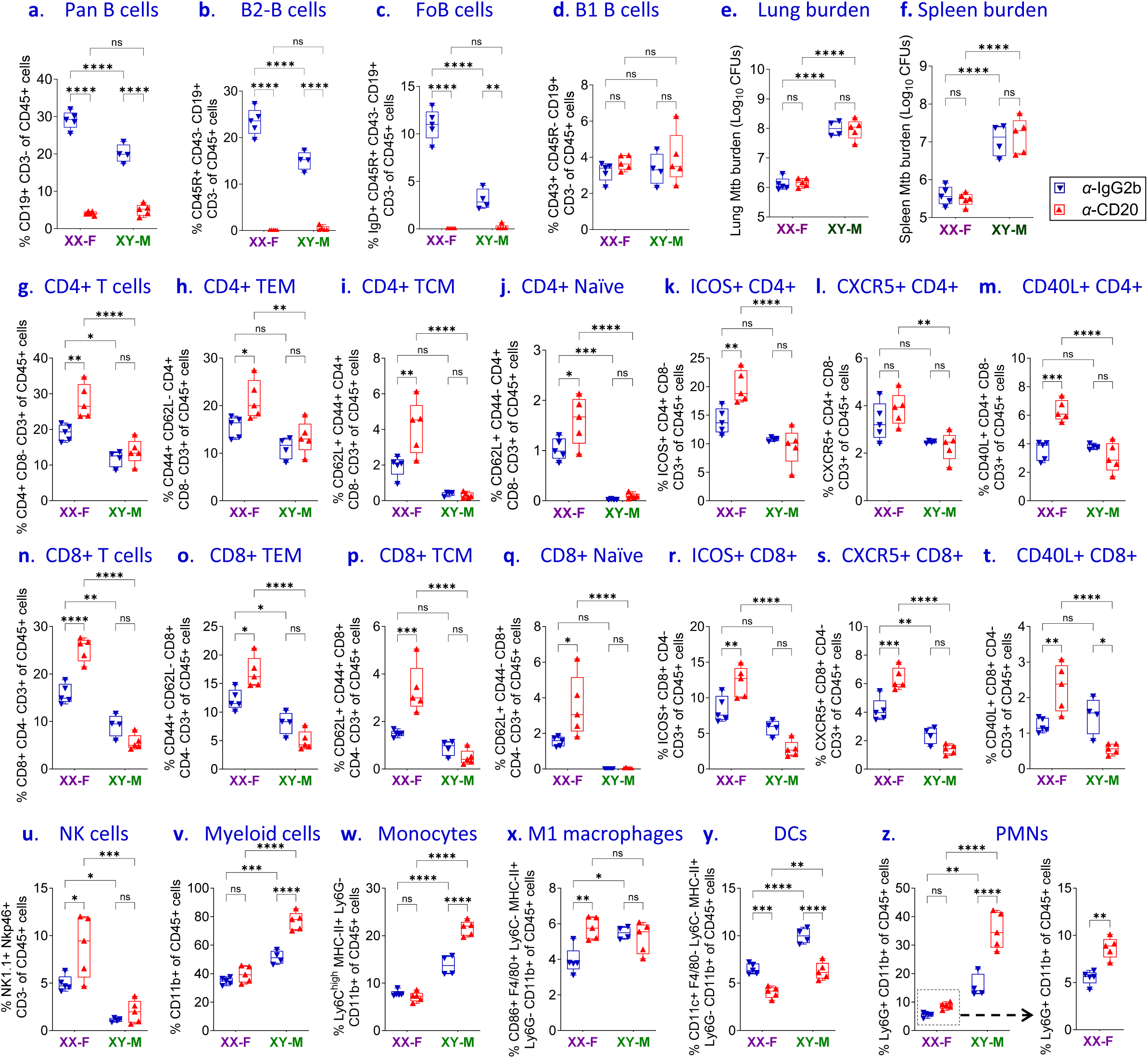
Loss of conventional B cells drives sex-specific spatial remodeling of the pulmonary immune microenvironment during chronic *Mtb* infection. Flow cytometric analysis of pulmonary leukocyte populations following anti-CD20 mAb-mediated depletion of conventional B cells in XXF and XYM mice, at 12 wpi with Mtb was performed. **(a),** Total B cells. **(b),** B2 B cells. **(c),** Follicular B (FoB) cells. **(d),** B1 B cells, depicting complete abrogation of B2 B cells. **(e),** Lung and **(f),** Spleen Mtb burden post-immunotherapy. Quantification of pulmonary CD4⁺ T-cell populations showing **(g),** total CD4⁺ T cells, **(h),** effector memory (TEM) CD4⁺ T cells, **(i),** central memory (TCM) CD4⁺ T cells, **(j),** naïve CD4⁺ T cells, **(k),** ICOS⁺ CD4⁺ T cells, **(l),** CXCR5⁺ CD4⁺ T cells, and **(m),** CD40L⁺ CD4⁺ T cells. Quantification of pulmonary CD8⁺ T-cell populations showing **(n),** total CD8⁺ T cells, **(o),** TEM CD8⁺ T cells, **(p),** TCM CD8⁺ T cells, **(q),** naïve CD8⁺ T cells, **(r),** ICOS⁺ CD8⁺ T cells, **(s),** CXCR5⁺ CD8⁺ T cells, and **(t),** CD40L⁺ CD8⁺ T cells. Quantification of innate immune cell populations showing **(u),** natural killer (NK) cells, **(v),** CD11b⁺ myeloid cells, **(w),** inflammatory monocytes, **(x),** M1 macrophages, **(y),** CD11c⁺CD11b⁺ dendritic cells (DCs), and **(z),** PMNs. Data are presented as box-and-whisker plots. n = 5 for all groups except for IgG2b isotype-control-treated XYM, where n=4, and each symbol represents an individual mouse. Data are plotted as mean ± SEM. Statistical significance was determined using two-way ANOVA with Tukey’s multiple comparisons test. *P < 0.05, **P < 0.01, ***P < 0.001, ****P < 0.0001, and ns, not significant.

Surprisingly, despite near-complete depletion of the B cell populations responsible for pulmonary follicle formation, anti-CD20 treatment had no measurable effect on bacterial burden in either the lungs or the spleens of XXF or XYM mice (Fig. 6e,f). Thus, although conventional B2/FoB B cells are indispensable for maintaining pulmonary immune spatial organization, they are not required for control of bacterial burden during established chronic TB in mice. Instead, these findings indicate that extensive remodeling of the pulmonary immune landscape may be the primary consequence of B cell depletion rather than altered antimicrobial immunity.

Remarkably, identical depletion of pulmonary B cells elicited fundamentally different immunological responses in females and males. As observed previously during chronic infection, isotype-treated XXF mice exhibited larger pulmonary CD4⁺ and CD8⁺ T cell compartments than XYM mice, whereas males displayed greater representation of myeloid populations, monocytes and dendritic cells, reflecting the sexually dimorphic immune landscape established during chronic TB. Following B cell depletion, however, immune remodeling diverged even further between the sexes.

In XXF mice, B cell depletion resulted in marked expansion of the adaptive immune compartment. Total CD4⁺ T cells increased significantly, together with both TEM and TCM subsets as well as naïve CD4⁺ T cells (Fig. 6g–j), suggesting broad enhancement of T cell differentiation and maintenance following the loss of organized B220+ lymphoid niches. Similarly, CD8⁺ T cells expanded substantially and were accompanied by increased frequencies of TEM, TCM and naïve CD8⁺ populations in B cell-depleted XXF mice (Fig. 6n–q). Importantly, both CD4⁺ and CD8⁺ T cells acquired a more activated phenotype characterized by increased ICOS, CXCR5 and CD40L expression (Fig. 6k–m,r–t), markers previously associated with protective T-cell responses and pulmonary lymphoid organization in this model. These findings indicate that pulmonary B cells are not required to sustain adaptive immunity in resistant females but instead function to spatially organize and regulate T cell activation within tertiary lymphoid structures during TB.

In striking contrast, XYM mice exhibited comparatively modest adaptive immune remodeling following B cell depletion. Rather than promoting extensive T cell expansion, loss of pulmonary B cells preferentially reshaped the innate immune compartment. In B cell-depleted XYM mice, total CD11b⁺ myeloid cells increased together with inflammatory monocytes (Fig. 6v,w), whereas M1 macrophage frequencies remained comparatively stable (Fig. 6x), indicating selective expansion of inflammatory rather than resident myeloid populations. Notably, although untreated XYM mice contained significantly higher frequencies of CD11c⁺CD11b⁺ dendritic cells than females during chronic infection, anti-CD20 treatment markedly reduced dendritic cell frequencies in both sexes (Fig. 6y), suggesting that maintenance of pulmonary dendritic cell networks is dependent upon intact B cell-rich lymphoid architecture irrespective of sex.

The most striking consequence of B cell depletion was observed within the neutrophil compartment. Although anti-CD20 treatment increased pulmonary PMN frequencies in both females and males, the response was dramatically amplified in XYM mice, resulting in robust neutrophilic accumulation that greatly exceeded that observed in females (Fig. 6z). This pronounced neutrophilia closely mirrored the elevated neutrophil-recruiting chemokines identified in Fig. 5, including CXCL1 and CXCL2, suggesting that pulmonary B cells normally restrain inflammatory neutrophil recruitment in susceptible males. Taken together, these findings demonstrate that pulmonary B cells orchestrate fundamentally distinct immune programs in female and male lungs during chronic TB. In resistant females, conventional B2 and follicular B cells maintain organized tertiary lymphoid structures that spatially regulate adaptive T cell activation and differentiation. Conversely, in susceptible males, where pulmonary BCFs are intrinsically sparse, endogenous B cells primarily suppress inflammatory myeloid remodeling, thereby limiting monocyte recruitment and, most notably, pathological neutrophil accumulation. Thus, disruption of pulmonary B cell organization redirects immunity toward adaptive immune expansion in females but excessive innate inflammatory responses in males, establishing a mechanistic link between loss of immune spatial organization, neutrophilia and the downstream NETotic pathology that develops during chronic TB.

### Loss of pulmonary B-2 B cell spatial organization promotes excessive NET formation during chronic TB

Having established that disruption of conventional B cell-dependent pulmonary organization preferentially promoted inflammatory myeloid remodeling and robust neutrophil accumulation in male mice (Figs. 5 and 6), we next asked whether these infiltrating neutrophils acquired a pathogenic phenotype characterized by NET formation. Lung sections were therefore examined by multiplex immunofluorescence using myeloperoxidase (MPO) and citrullinated histone H3 (H3Cit), a canonical marker of NETosis (Fig. 7a). Consistent with the flow cytometric analyses, anti-CD20 treatment increased pulmonary MPO⁺ neutrophil infiltration in both XXF and XYM mice, although the magnitude of this response was substantially greater in males (Fig. 7a,b). Histological examination further revealed abundant extracellular H3Cit-positive structures throughout anti-CD20-treated lungs, particularly within inflammatory lesions of XYM mice, whereas comparatively fewer NETotic structures were detected in females (Fig. 7a). Quantitative analysis confirmed a significant increase in both the number of H3Cit-positive foci and the overall H3Cit-positive tissue area following B-cell depletion in XYM but not XXF mice (Fig. 7c,d). Collectively, these findings establish NETosis as a downstream consequence of B-cell depletion-induced inflammatory remodeling during TB. Whereas disruption of conventional B-cell-rich pulmonary niches in females primarily reshaped adaptive immune responses with relatively limited pathological consequences, the same perturbation in males unleashed excessive neutrophil recruitment culminating in widespread NET formation.

**Figure 7.**
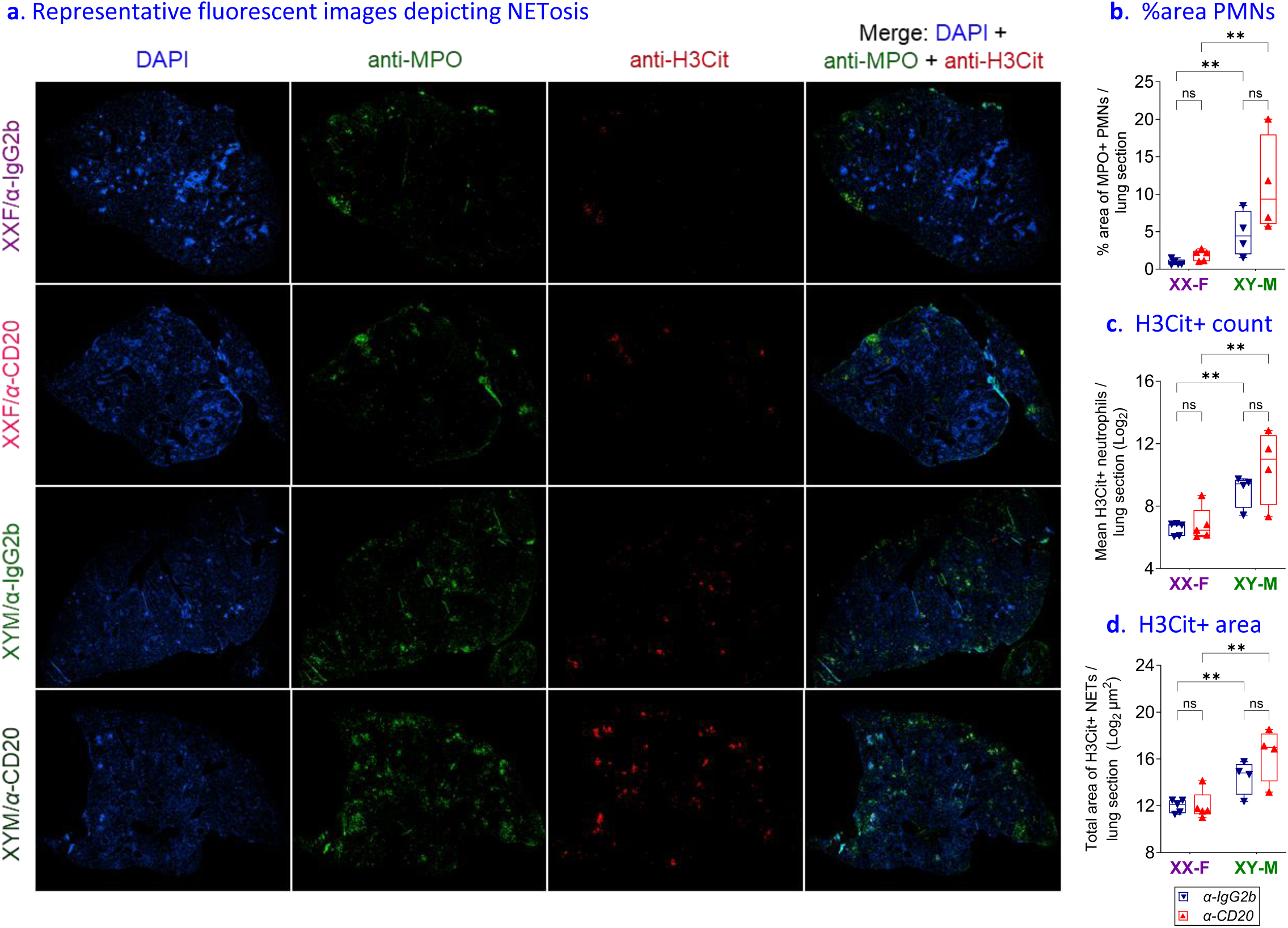
Conventional B-cell depletion promotes exaggerated neutrophil extracellular trap formation in male lungs during chronic *Mtb* infection. **(a),** Multiplex immunofluorescence and analyses of NET formation following anti-CD20 mAb-mediated depletion XXF and XYM mice at 12 wpi. Representative multiplex immunofluorescence images of lung sections stained with DAPI (blue), myeloperoxidase (MPO; green), and citrullinated histone H3 (H3Cit; red), with merged images depicting MPO⁺H3Cit⁺ neutrophil extracellular traps. Quantification of **(b),** % MPO⁺/green PMN area per lung section, **(c),** H3Cit⁺ cell counts per lung section, and **(d),** Total H3Cit⁺ tissue area per lung section. Data are presented as box-and-whisker plots showing individual mice. n = 4 mice per group, and each symbol represents an individual mouse. Data are plotted as mean ± SEM. Statistical significance was determined using two-way ANOVA with Tukey’s multiple comparisons test. *P < 0.05, **P < 0.01, ***P < 0.001, ****P < 0.0001, and ns, not significant.

## Discussion

Sex differences are a hallmark of TB, yet the mechanisms through which biological sex shapes protective immunity remain poorly understood^4^. Using the FCG model together with adoptive transfer, pathway-specific blockade, and B cell depletion, we demonstrate that sex-dependent immunity during chronic Mtb infection is governed by two complementary mechanisms. First, female-derived CD4⁺ and CD8^+^ T cells possess an intrinsically superior protective program that enhances bacterial control independently of sex chromosome complement. Second, pulmonary B cells regulate the spatial organization of immune responses, coordinating adaptive immunity in females while restraining pathological inflammatory remodeling in males. Together, these findings identify cell-intrinsic immune programming and tissue-level immune organization as complementary determinants of sexually dimorphic immunity during chronic TB.

Our adoptive transfer studies establish that the enhanced resistance observed in females is primarily attributable to the CD4⁺ T cell compartment. Female-derived, but not XX male-derived, CD4⁺ T cells conferred protection to susceptible male recipients, reducing pulmonary and splenic bacterial burden, increasing pulmonary Bcl6⁺ CD4⁺ T cells, and limiting neutrophilic inflammation. Because donor cells shared an identical XX chromosome complement, these findings identify gonadal sex as the dominant determinant of protective CD4⁺ T cell programming. Although sex hormones are known to influence T cell differentiation^28^, our data demonstrate that these intrinsic differences persist during chronic infection and are sufficient to reshape the immune landscape of a susceptible host.

Effective control of TB depends not only on antimicrobial immunity but also on coordinated regulation of inflammation^29, 30^. Female-derived CD4⁺ T cell transfer simultaneously enhanced IFN-γ production while suppressing inflammatory cytokines and neutrophil-recruiting chemokines, resulting in reduced pulmonary pathology despite improved bacterial control. These findings suggest that female adaptive immunity promotes a more balanced immune response rather than simply amplifying inflammation. More broadly, our data support a model in which the granulomatous microenvironment is a critical regulator of disease outcome^31, 32^. In susceptible males, failure to establish an organized adaptive immune niche favors uncontrolled myelopoiesis, hyperinflammatory neutrophilic responses and poor coordination of T and B cell immunity, ultimately accelerating disease progression.

Mechanistically, we identify CXCR3- and CD40L-dependent pathways as key regulators of female-associated protection against TB. Blockade of either pathway impaired bacterial containment, demonstrating that both pathways contribute to protective immunity during chronic Mtb infection. Whereas CXCR3 inhibition predominantly altered activated and memory T-cell populations, CD40L blockade profoundly disrupted pulmonary B cell organization, highlighting complementary roles in coordinating adaptive immunity and lymphoid architecture. Interestingly, increased bacterial burdens following pathway blockades were accompanied by reduced histopathology scores, reflecting the loss of organized B cell-rich lymphoid structures rather than diminished inflammation. These findings emphasize that conventional histopathological assessment does not necessarily distinguish protective immune organization from destructive inflammatory pathology.

A major finding of this study is that pulmonary B cells function primarily as organizers of immune spatial architecture rather than direct mediators of bacterial control^15, 33, 34^. Depletion of conventional B2 and follicular B cells abolished pulmonary BCFs in both sexes but had no measurable effect on bacterial burden, indicating that these cells are dispensable for controlling established chronic TB despite the simultaneous loss of their antigen-presenting and immunomodulatory functions. The preservation of bacterial containment likely reflects functional redundancy among professional antigen-presenting cells, including dendritic cells, macrophages, and neutrophils, which can compensate for the absence of B cell-mediated antigen presentation and sustain protective T cell responses. Rather than impairing bacterial control, B cell depletion profoundly reshaped the pulmonary immune landscape in a sex-dependent manner. In females, disruption of BCFs promoted expansion and activation of both CD4⁺ and CD8⁺ T cells, consistent with a role for organized B cell niches in coordinating local adaptive immunity rather than simply increasing T cell abundance. In contrast, males exhibited comparatively modest adaptive immune changes but marked inflammatory remodeling characterized by increased inflammatory monocytes, elevated neutrophil-attracting chemokines, pronounced neutrophilia and enhanced NET formation. NETs are abundant within necrotic granulomas of TB patients and non-human primates and have been implicated in promoting Mtb persistence^35^, however, their clear role in Mtb clearance is still controversial and being investigated^36, 37^. In this milieu, the increased NETosis following B cell depletion identifies loss of pulmonary immune organization as a potential upstream driver of pathological neutrophil activation^38^. However, despite enhanced NET formation, bacterial burden remained unchanged in both males and females, suggesting that NET-associated pathology can be uncoupled from bacterial control during chronic infection^37, 39, 40^. Together, these findings demonstrate that pulmonary B cells primarily regulate immune organization and inflammatory homeostasis, with fundamentally distinct functions in the two sexes—promoting coordinated adaptive immunity in females while restraining excessive innate inflammation in males.

The role of pulmonary BCFs in TB remains controversial. Although accumulating evidence supports protective functions for B cells and antibodies during Mtb infection, B cell-deficient mouse models have generally reported little or no effect on pulmonary bacterial burden^19, 41^. Our findings extend these observations by showing that elimination of pulmonary BCFs similarly does not compromise bacterial control, but instead triggers extensive remodeling of local immune networks. Whether these follicles actively contribute to host protection or represent ectopic “germinal center-like’ structures generated through lymphoid neogenesis during chronic inflammation remains unresolved^42^. Nevertheless, the marked immunological changes that follow their disruption strongly argue against pulmonary BCFs being passive bystanders and instead support their role as active organizers of the local immune microenvironment, where they coordinate immune cell positioning, cellular interactions and inflammatory responses within the infected lung^15, 43, 44^.

Collectively, our findings support a model in which biological sex determines the outcome of chronic TB through two interconnected layers of immune regulation. Intrinsic programming of female CD4⁺ T cells promotes effective bacterial control while limiting inflammatory pathology, whereas pulmonary B cells establish the spatial organization required to coordinate adaptive immunity in females and suppress pathological neutrophilic remodeling in males. These findings extend the concept of sex-biased immunity beyond differences in immune cell composition to include the organization of immune responses within infected tissues. Therapeutic strategies aimed at preserving protective immune architecture or limiting pathological neutrophilic remodeling may therefore complement conventional antimicrobial therapy and provide a framework for developing sex-informed host-directed therapies for TB.

## Acknowledgements

We gratefully acknowledge support from the National Institutes of Health grants R37AI167750 and P30AI16843. We thank the SKCCC OTIS Core for assistance with histology and immunohistochemistry. Schematic figures and the graphical abstract were created using BioRender (Academic License). We also acknowledge past and present members of the Bishai laboratory for their valuable discussions and suggestions throughout the study.

## Materials and methods

### Ethics statement

All animal procedures were performed under protocol M022M466, which was approved by the Institutional Animal Care and Use Committee (IACUC) at the School of Medicine, Johns Hopkins University (JHSOM). Animals were maintained in individually ventilated cages, with a maximum of five mice of the same strain housed together. Standardized husbandry conditions included a 12-hour light/dark cycle, room temperatures maintained between 20 and 24 °C, relative humidity of 45–65%, and unrestricted access to food and drinking water. Upon completion of the study, mice were euthanized in accordance with IACUC-approved procedures to ensure humane treatment and minimize discomfort.

### Mice breeding and genotyping

Breeder XY⁻ males for the Four Core Genotypes (FCG) mouse model were provided by Arthur P. Arnold to Sabra L. Klein^26^. FCG colonies were established by crossing XY⁻ males with wild-type C57BL/6J females (Strain #000664, Jackson Laboratory), generating four genotypes: XX females (XXF), XY females (XYF), *Sry*-harboring XX males (XXM), and XY males (XYM). At weaning, mice were separated by gonadal sex, and genotypes were confirmed by triplex PCR for *Sry* and the Y-chromosome marker *Ssty*, as previously described^45^. Littermates of the same genotype were co-housed and used for infection at 8 weeks of age. FCG mice carry the endogenous C57BL/6J *Ptprc^b^* (CD45.2/Ly5.2) allele. For adoptive transfer studies, female JAXBoy mice expressing the *Ptprc^a^* (CD45.1/Ly5.1) allele were purchased from the Jackson Laboratory and maintained under specific pathogen-free conditions, as previously described.

### Bacterial strain and aerosol infection

The *Mycobacterium tuberculosis* H37Rv strain was obtained from the Johns Hopkins Center for Tuberculosis Research and cultured in Middlebrook 7H9 medium supplemented with 10% (v/v) OADC, 0.5% (v/v) glycerol, and 0.05% (v/v) Tween 80. Mid-log phase cultures (OD₆₀₀ ≈ 1.0) were aliquoted (1 mL) and stored at −80 °C.

For all the infection studies, mice were housed five per cage under biosafety level 3 (BSL-3) conditions at the JHSOM animal facility, with unrestricted access to food and water. Animals aged 8–9 weeks were infected via aerosol exposure using a Glas-Col Inhalation Exposure System (Terre Haute, IN). Freshly thawed bacterial stocks were diluted in sterile phosphate-buffered saline (PBS, pH 7.4) to achieve the target inoculum based on prior calibration. All the mouse groups were infected in parallel to minimize batch-to-batch variation. At 24 h post-infection, 4–5 mice per run were euthanized to confirm implantation by lung CFU enumeration. Mice were monitored weekly for body weight and clinical status. All infectious work was performed in BSL-3 containment, and moribund animals were humanely euthanized.

### Adoptive transfer

To determine whether gonadal sex intrinsically programs adaptive immune cells to differentially regulate host resistance to TB, adoptive transfer experiments were performed. Congenic XX female (XX-F; CD45.1) and XX male (XX-M; CD45.2) mice were used as donors, whereas chronically infected XY male (XY-M; CD45.2) mice served as recipients (Fig. 1a). Donor and recipient mice were infected simultaneously with approximately 100 CFUs of Mtb H37Rv as described above. Four weeks post-infection, donor mice were euthanized, and lungs were harvested under sterile conditions. Lung tissues were dissociated in gentleMACS C Tubes (Miltenyi Biotec) containing Mouse Lung Dissociation Kit enzymes (Cat. No. 130-095-927) using a gentleMACS Dissociator with the 37C_m_LDK_1 program. Cell suspensions were filtered through 70-μm strainers, washed, resuspended in IMDM containing 10% FBS, and viable cells were quantified by trypan blue exclusion using a Countess Automated Cell Counter (Thermo Fisher Scientific). Pulmonary CD4⁺ T cells and pan-B cells were purified by magnetic negative selection using the MojoSort™ Mouse CD4 T Cell Isolation Kit (BioLegend, Cat. No. 480033) and the MojoSort™ Mouse Pan B Cell Isolation Kit (BioLegend, Cat. No. 480052), according to the manufacturer’s instructions. Purified CD4⁺ T cells or CD19⁺ B cells were resuspended in sterile PBS pH 7.4, and 2 × 10⁶ cells were administered via the intraperitoneal route into chronically infected XY-M (CD45.2) recipient mice. Recipient mice received one of four donor cell populations: XX-F-derived CD4⁺ T cells, XX-M-derived CD4⁺ T cells, XX-F-derived CD19⁺ B cells, or XX-M-derived CD19⁺ B cells. Cell transfers were initiated at 4 wpi and repeated weekly until week 11 post-infection to maintain donor lymphocyte engraftment throughout chronic disease. Recipient mice were euthanized at 12 wpi, and lungs and spleens were collected for determination of bacterial burden by CFU enumeration, histopathological analysis, multiplex immunofluorescence staining, cytokine measurements, and flow cytometric immunophenotyping. Donor-derived cells were distinguished from recipient leukocytes using congenic CD45 allelic markers (CD45.1 and CD45.2), enabling assessment of donor cell engraftment following adoptive transfer.

### Antibody-mediated immunotherapy

Mice were infected with a low-dose aerosol of Mtb H37Rv. To interrogate the role of CD40L- and CXCR3-dependent immune pathways during established infection, treatment was initiated at 4 wpi, coinciding with the onset of adaptive immunity (Fig. 3E). Mice received intraperitoneal injections of Ultra-LEAF™ purified anti-CD154 (CD40L; clone MR1) or Ultra-LEAF™ purified anti-CXCR3 (clone CXCR3-173) monoclonal antibodies (BioLegend) at 250 μg/mouse twice weekly until 12 wpi. Control animals received an equivalent dose of Armenian hamster IgG isotype control antibody (clone HTK888) on the same schedule. At 12 wpi, lungs were harvested for bacterial burden determination, histopathological analysis, cytokine profiling, and flow-cytometric immunophenotyping.

To determine the contribution of B cells during chronic TB, mice were infected as described above and treated with a depleting Ultra-LEAF™ purified anti-CD20 monoclonal antibody (clone SA271G2; BioLegend) at 250 μg per mouse by intraperitoneal injection once weekly during the chronic phase of infection, as indicated in the experimental schematic (Fig. 5A). Control animals received a matched dose of rat IgG2b isotype control antibody (clone RTK4530). Both XXF and XYM mice were included in these experiments. At the experimental endpoint, lungs were collected for bacterial burden determination, histopathological analysis, and flowcytometric immunophenotyping.

### Organ collection and enumeration of Mtb burden

At pre-defined post-infection time points, animals were euthanized and final body mass was documented. Whole blood was obtained using BD Microtainer serum tubes, followed by sterile excision and weighing of lung and spleen tissues. Lungs were divided for downstream applications: right lobes were allocated for Mtb load assessment, RNA isolation, and cytokine/chemokine analyses, whereas the left lobe was preserved in 10% neutral-buffered formalin for histological examination and immunostaining. Weights of intact lungs and individual lobar fractions were recorded to enable total burden calculations.

For CFU analysis, right lung lobes and spleens were homogenized in 2.5 mL sterile PBS using Precellys^®^ 7 mL Tissue Grinding bead-beating tubes (CKMix50). Tissue suspensions were serially diluted, and 0.5 mL volumes were plated onto Middlebrook 7H11 agar (Difco) containing 10% OADC, 0.5% glycerol, and antimicrobial supplements (cycloheximide 10 mg/mL, carbenicillin 50 mg/mL, polymyxin B 25 mg/mL, and trimethoprim 20 mg/mL; Sigma-Aldrich). Cultures were incubated at 37 °C for 3–4 weeks prior to colony enumeration. Bacterial counts were normalized for dilution factors, plated volume, and lung partitioning and reported as log₁₀ CFU per organ.

### Pulmonary cytokine and chemokine profiling

Pulmonary cytokine and chemokine levels were measured at pre-defined post-infection time points using mouse LEGENDplex multiplex assays (BioLegend). Briefly, frozen lung samples were mechanically disrupted by bead homogenization in 2 mL Precellys CK28-R tubes (Bertin Corp.) containing PBS supplemented with protease inhibitors (Halt™, Thermo Fisher Scientific). Lysates were centrifuged at 10,000 rpm for 10 min at 4 °C, and clarified supernatants were passed through 0.2 μm filters. Aliquots were stored at −80 °C following snap-freezing in liquid nitrogen until analysis. Cytokines were quantified using the LEGENDplex Mouse Inflammation Panel (13-plex; Cat. #740150), while chemokines were assessed using the Mouse Proinflammatory Chemokine Panel 1 (13-plex; Cat. #741294) and Panel 2 (8-plex; Cat. #741067) according to the manufacturer’s instructions. Acquisition was performed on a CytoFLEX flow cytometer (Beckman Coulter), and concentrations were calculated using LEGENDplex Qognit software (BioLegend). All analyte levels were normalized to total protein content determined by Pierce™ BCA Protein Assay Kits (Thermo Fisher Scientific).

### Histopathological analysis

The left pulmonary lobe from each animal was preserved in 10% neutral-buffered formalin for 72 h before routine paraffin processing. Tissue blocks were sectioned at 4 µm, stained with hematoxylin and eosin (H&E), and prepared by the Johns Hopkins University Oncology Tissue and Imaging Service (OTIS) Core. Bright-field whole-slide images were acquired at ×40 magnification using a Hamamatsu Nanozoomer S210 digital pathology scanner (Hamamatsu Photonics, Shizuoka, Japan) and imported into Concentric LS for Research (v4.4; Proscia Inc.) for analysis. Histopathological scoring and lesion quantification were performed in a blinded manner by an American College of Veterinary Pathologists (ACVP)-boarded veterinary pathologist. Semi-quantitative histopathology scores were assigned to each specimen based on the severity and extent of pulmonary injury and inflammation, including lung consolidation, accumulation of foamy alveolar macrophages, perivascular leukocyte cuffing, peribronchiolar leukocyte cuffing, neutrophilic infiltration, and necrosis. Scoring was performed according to the criteria outlined in Supplementary Table S1. Tissues considered within normal limits were assigned a score of 0. Data visualization and statistical analyses were performed using GraphPad Prism.

### Immunohistochemical analysis

Immunohistochemistry was performed on serial 4-µm sections prepared from formalin-fixed, paraffin-embedded mice lungs to detect B lymphocytes using an anti-CD45R/B220 antibody. All staining was completed by the Sidney Kimmel Comprehensive Cancer Center Oncology Tissue and Imaging Service (OTIS) Core at Johns Hopkins University (Baltimore, MD) using a Ventana Discovery Ultra automated staining platform (Roche Diagnostics). Following automated deparaffinization and hydration, antigen retrieval was carried out with Ventana Ultra CC1 buffer (Roche Diagnostics; Cat. #6414575001) at 96 °C for 64 min. Sections were incubated with rat anti-mouse CD45R/B220 antibody (clone RA3-6B2; 1:300; BD Transduction Laboratories; Cat. #3088544) at 36 °C for 40–60 min. Because the primary antibody was rat-derived, signal amplification was achieved with a rabbit anti-rat linker antibody (1:500; Vector Laboratories; Cat. #AI4001) for 32 min at 36 °C. Bound antibody was detected using the anti-rabbit HQ detection system (Roche Diagnostics; Cat. ##7017936001 and #7017812001), followed by visualization with the ChromoMap DAB detection kit (Roche Diagnostics; Cat. #5266645001). Slides were counterstained with Mayer’s hematoxylin, dehydrated, and coverslipped using routine procedures.

Brightfield whole-slide images were acquired at ×40 magnification (0.23 µm/pixel) using a Hamamatsu Nanozoomer S210 scanner (Hamamatsu Photonics, Shizuoka, Japan). Digital images were analyzed on the Concentriq pathology platform (Proscia, Philadelphia, PA) with identical display settings for all specimens. For assessment of lung consolidation, total lung area and consolidated regions were manually annotated in QuPath, and the percentage of consolidated lung area was calculated relative to the total lung area. For immunohistochemical analysis, DAB-positive staining was quantified in QuPath using the “Estimate Stain Vectors” function for color deconvolution, followed by stain thresholding and positive-cell detection to determine the number of positively stained cells. Identical threshold values and detection parameters were applied across all tissue sections to ensure consistency in analysis^46^.

### Multiplex confocal immunofluorescence

Formalin-fixed, paraffin-embedded lung specimens were sectioned (4 µm), mounted on Superfrost Plus slides, and processed through deparaffinization, rehydration, and heat-mediated antigen retrieval in citrate buffer (Abcam) for 30 min. Autofluorescence was suppressed using TrueBlack® (Biotium), followed by blocking with 5% normal goat serum for 1 h at room temperature. Sections were incubated overnight at 4 °C with anti-myeloperoxidase (MPO; AF3667, R&D Systems; 1:100) and anti-citrullinated histone H3 (H3Cit; clone EPR20358-120, Abcam; 1:100). Species-specific Alexa Fluor 488- and 555-conjugated secondary antibodies (Invitrogen; 1:1000) were applied for 1 h, and nuclei were labeled with DAPI (1 µg/mL). Images were acquired using a Nikon Eclipse Ti2-E confocal microscope under identical acquisition settings for all samples. Fluorescence quantification was performed in ImageJ/Fiji using an identical analysis workflow for all samples. Following the application of uniform threshold settings, raw integrated fluorescence intensity was determined across the entire tile-scanned lung section. Lung specimens from four mice per group were analyzed, and each section contributed a single value for subsequent statistical analysis.

### Lung dissociation and multicolor flow cytometry

Right lung lobes were collected into gentleMACS C Tubes (Miltenyi Biotec) containing enzymes from the Mouse Lung Dissociation Kit (Miltenyi Biotec; Cat. #130-095-927) and processed using a gentleMACS Dissociator with the 37C_m_LDK_1 program. Cell suspensions were filtered through 70-µm cell strainers, washed, subjected to erythrocyte lysis using ACK buffer, and resuspended in IMDM supplemented with 10% FBS. Viable cells were enumerated by trypan blue exclusion using a Countess automated cell counter (Thermo Fisher Scientific). For flow cytometric analysis, 2 × 10^6^ viable cells were stained per sample. Cells were first labeled with Zombie Red™ viability dye (BioLegend), followed by Fc receptor blockade using TruStain FcX™ PLUS (BioLegend), and subsequently stained with optimized panels of fluorochrome-conjugated antibodies. Intracellular transcription factor staining was performed using the FOXP3 Fix/Perm Buffer Set (BioLegend) according to the manufacturer’s instructions. Following staining, cells were washed, resuspended in Cyto-Last™ Buffer (BioLegend), filtered through strainer-cap FACS tubes, and acquired on BD LSR II or BD LSRFortessa flow cytometers. Data were analyzed using FlowJo v10.10 (Tree Star). Immune cell populations were identified using the following fluorochrome-conjugated antibodies: Brilliant Violet 510™ anti-mouse CD45, Pacific Blue™ anti-mouse CD3, PerCP anti-mouse CD4, Alexa Fluor® 700 anti-mouse CD8a, Brilliant Violet 605™ anti-mouse CD19, KIRAVIA Blue 520™ anti-mouse NK-1.1, PE/Cyanine7 anti-mouse CD335 (NKp46), Alexa Fluor® 647 anti-mouse/human Bcl-6, Brilliant Violet 510™ anti-mouse CD45.1, Alexa Fluor® 700 anti-mouse CD11b, APC/Fire™ 750 anti-mouse F4/80, Brilliant Violet 650™ anti-mouse CD86, BUV563 rat anti-mouse Ly-6G, Brilliant Violet 785™ anti-mouse Ly-6C, KIRAVIA Blue 520™ anti-mouse CD11c, Spark UV387™ anti-mouse I-A/I-E, PerCP/Cy5.5 anti-mouse CD3, APC/Fire™ 750 anti-mouse CD4, KIRAVIA Blue 520™ anti-mouse CD62L, Brilliant Violet 650™ anti-mouse CD44, Alexa Fluor® 700 anti-mouse CD3, Brilliant Violet 421™ anti-mouse CD45R/B220, APC/Fire™ 750 anti-mouse CD19, Alexa Fluor® 488 anti-mouse CD5, APC anti-mouse CD43, Brilliant Violet 650™ anti-mouse CD138, PE/Cyanine7 anti-mouse/rat/human CD27, PerCP/Cyanine5.5 anti-mouse IgD, Brilliant Violet 605™ anti-mouse CD45R, APC anti-mouse ICOS, FITC anti-mouse CXCR3, PE/Cyanine7 anti-mouse CD185 (CXCR5), and PE anti-mouse CD40L. Antibody concentrations were optimized prior to use. Compensation was performed using single-stained controls, and gating boundaries were established using fluorescence-minus-one (FMO) controls generated from lung cell suspensions. Representative gating strategies are provided in the Supplementary Figures.

### RNA isolation and quantitative real-time PCR

Total RNA was extracted from flash-frozen lungs of Mtb-infected FCG mice by homogenization in QIAzol followed by purification with the miRNeasy Mini Kit (Qiagen). Genomic DNA contamination was eliminated by DNase I treatment (Promega), and first-strand cDNA was generated using the High-Capacity cDNA Reverse Transcription Kit (Thermo Fisher Scientific). Quantitative PCR was carried out with PowerUp™ SYBR Green Master Mix (Thermo Fisher Scientific; A25742) on StepOne Plus or QuantStudio 3 Real-Time PCR instruments (Applied Biosystems). Primers were designed using the GenScript Real-Time PCR Primer Design Tool and synthesized by Integrated DNA Technologies. The following primer pairs were used: *Cxcr3*, forward 5′-CTACGATCAGCGCCTCAATG-3′ and reverse 5′-TCTGGAGACCAGCAGAACAG-3′; *Cd40lg*, forward 5′-CTAATCGGGAGCCTTCGAGT-3′ and reverse 5′-CCCAAGTGAACAGACTGCTG-3′. Amplification data were processed with the corresponding Applied Biosystems software. Gene expression was normalized to 18S rRNA, and relative transcript abundance was calculated using the 2^−ΔΔCt^ method. ΔΔCt values were referenced to the mean ΔCt of α-IgG2b-XXF as controls, and results are presented as fold change.

### Statistical analysis

Statistical analyses were performed using GraphPad Prism (v10; GraphPad Software). Depending on the experimental design, data were analyzed using one- or two-way analysis of variance (ANOVA), followed by Tukey’s multiple comparisons test. Statistical tests used for individual experiments are specified in the corresponding figure legends. CFU data were log₁₀-transformed before analysis. Results are presented as mean ± SEM, and a two-sided P < 0.05 was considered statistically significant. Sample sizes were not predetermined by statistical methods. Mice were randomly assigned to experimental groups after aerosol infection, and all collected data were included in the analyses. Occasional mortality during infection reduced sample sizes in selected cohorts. Investigators were not blinded to group allocation during experimentation or data analysis, and no data were excluded.

---

**Supplementary Table S1.** Histopathology scoring.

Histopathology scoring
| Category | Criteria | Score |
| --- | --- | --- |
| Severity of lung injury | <5% area involvement | 1 |
|  | 6-10% area involvement | 2 |
|  | 10-20% area involvement | 3 |
|  | 21-50% area involvement | 4 |
|  | >50% area involvement | 5 |
| Number of foamy alveolar macrophages | <3 cell aggregates | 1 |
|  | 4-9 cell aggregates | 2 |
|  | >10 cell aggregates | 3 |
| Number of perivascular leukocyte cuffings | <3 cell aggregates | 1 |
|  | <4-9 cell aggregates | 2 |
|  | >10 cell aggregates | 3 |
| Number of peribronchiolar leukocyte cuffings | <3 cell aggregates | 1 |
|  | 4-9 cell aggregates | 2 |
|  | >10 cell aggregates | 3 |
| Number of neutrophils | <3 cell aggregates | 1 |
|  | 4-9 cell aggregates | 2 |
|  | >10 cell aggregates | 3 |
| Necrosis | Rare | 1 |
|  | Scattered | 2 |
|  | Prominent | 3 |

